# Partitioning the colonization and extinction components of beta diversity: Spatiotemporal species turnover across disturbance gradients

**DOI:** 10.1101/2020.02.24.956631

**Authors:** Shinichi Tatsumi, Joachim Strengbom, Mihails Čugunovs, Jari Kouki

## Abstract

Changes in species diversity often result from species losses and gains. The dynamic nature of beta diversity (i.e., spatial variation in species composition) that derives from such temporal species turnover, however, has been largely overlooked. Here, we disentangled extinction and colonization components of beta diversity by using the sets of species that went locally extinct and that newly colonized the given sites. We applied this concept of extinction and colonization beta diversity to plant communities that have been repeatedly measured in experimentally disturbed forests. We first found no difference in beta diversity across disturbance gradients when it was analyzed for communities at a single point in time. From this result, we might conclude that disturbance caused no impact on how species assemble across space. However, when we analyzed the extinction and colonization beta diversity, both measures were found to be significantly lower in disturbed sites compared to undisturbed sites. These results indicate that disturbance removed similar subsets of species across space, making communities differentiate, but at the same time induced spatially uniform colonization of new species, causing communities to homogenize. Consequently, the effects of these two processes canceled each other out. The relative importance of extinction and colonization components *per se* also changed temporally after disturbance. Analyses using extinction and colonization beta diversity allowed us to detect nonrandom dis- and re-assembly dynamics in plant communities. Our results suggest that common practices of analyzing beta diversity at one point in time can mask significant variation driven by disturbance. Acknowledging the extinction–colonization dynamics behind beta diversity is essential for understanding the spatiotemporal organization of biodiversity.

## INTRODUCTION

Ecological disturbance initiates community assembly where species dynamically turnover (Connell 1978, Caswell and Cohen 1991, Fukami and Nakajima 2011). With recent rises in the severity and frequency of disturbance worldwide (Nyström et al. 2000, Seidl et al. 2017), disturbance-induced reorganizations of species assemblages have become increasingly relevant in ecosystem management and conservation (Mori 2011). Despite substantial efforts, however, links between disturbance and community assembly are not yet generalizable (Jiang and Patel 2008, Myers et al. 2015, Tatsumi et al. 2019). In particular, compared with local and regional species richness (i.e., alpha and gamma diversity, respectively), little is known about how the spatial variation in species composition (i.e., beta diversity) is influenced by the temporal species turnover associated with disturbance (Vellend et al. 2007, Jiang and Patel 2008, Fukami and Nakajima 2011). Decreases in beta diversity, often referred to as biotic homogenization, can impair ecosystem functionality at large spatial scales (van der Plas et al. 2016, Hautier et al. 2018, Mori et al. 2018) and have been observed in disturbed landscapes (Vellend et al. 2007). A deeper understanding of the dynamic processes behind beta diversity would provide a critical step towards predicting how the landscape-scale organization of biodiversity and ecosystem functionality might change in response to future disturbance regimes.

Disturbance-induced community assembly should consist of two components: species losses (i.e., local extinctions) and gains (i.e., colonization). To our knowledge, however, no attempt has yet been made to explicitly disentangle these two components in shaping post-disturbance beta diversity. As such, we defined six cases that describe how local extinction and colonization drive community homogenization and differentiation (Fig. 1). First, extinction can decrease beta diversity when non-widespread species (e.g., habitat specialists) are selectively excluded from the entire region through direct physical damage or environmental alterations (Henle et al. 2004; Myers et al. 2015) (case 1 in Fig. 1). Conversely, extinction can increase beta diversity by excluding a similar set of species (Attum et al. 2006) (case 2) or different species across local patches (Cutler 1991) (case 3). Colonization can increase beta diversity if disturbance promotes spatial variation in population establishment by increasing environmental heterogeneity among sites (Myers et al. 2015) or if it selects for species with low to moderate dispersal abilities that would result in constrained levels of dispersal among local patches (Mouquet and Loreau 2003, Chase 2007) (case 4). In contrast, colonization can reduce beta diversity if disturbance homogenizes the environment across space under the condition of ample external dispersal from the species pool (Catano et al. 2017) (case 5). Furthermore, limited colonization rates of specialists can decrease beta diversity by resulting in widespread occurrence of a similar suite of generalists irrespective of local habitat conditions (Vellend et al. 2007) (case 6). Note that beta diversity can reflect two different phenomena (namely, species turnover and nestedness; Baselga 2010), but we do not distinguish between them here for simplicity. The recently-developed concept of temporal beta diversity (Legendre and Gauthier 2014, Legendre 2019) also differs from what we propose here (Fig. 1), fundamentally because while the temporal beta diversity directly compares community composition between two different times, we focus here on the subsets of community members that were lost and gained.

**Fig. 1.**
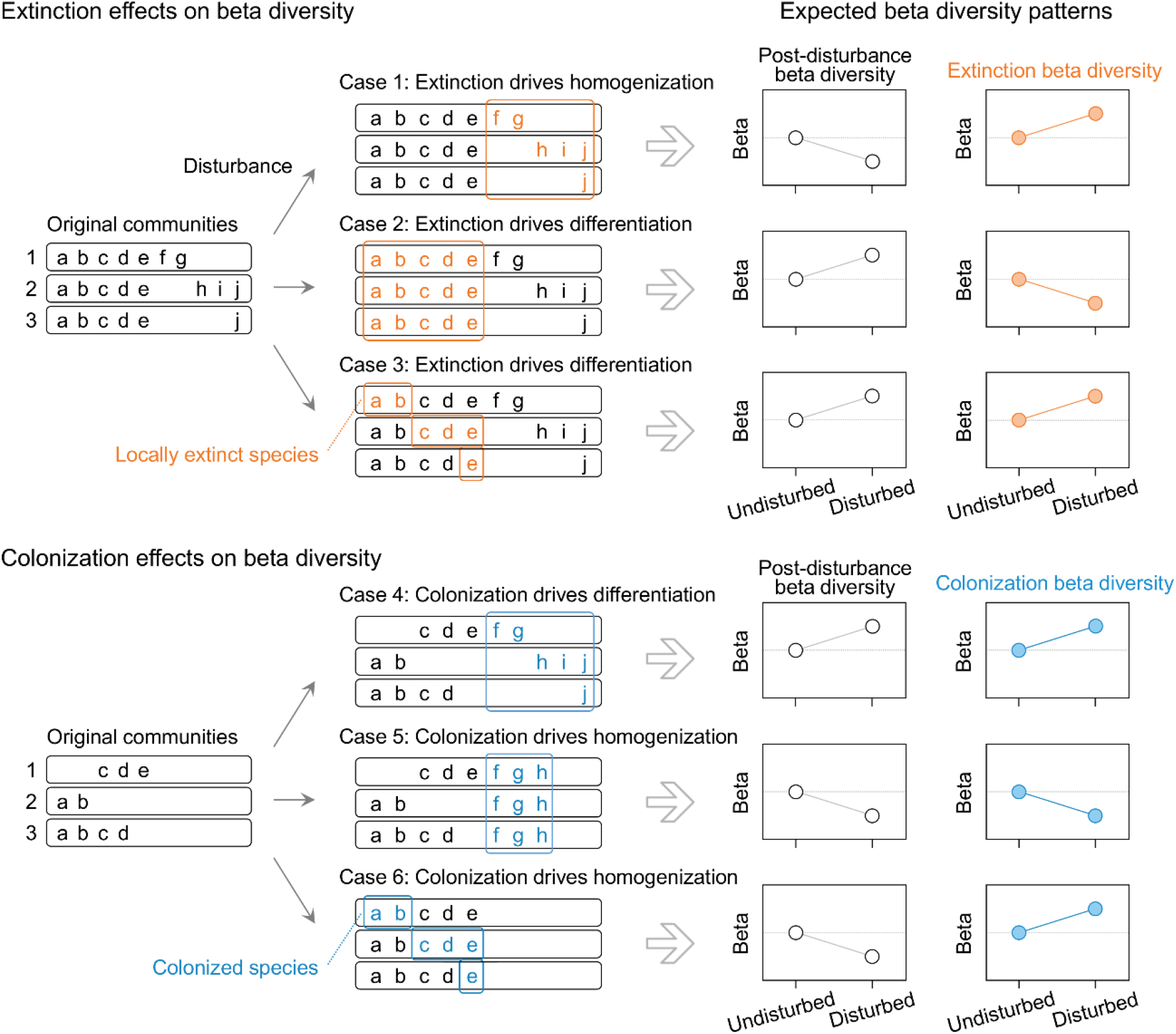
Schematic representation of the links between disturbance-induced extinction, colonization, and the resultant patterns of beta diversity. Different letters indicate different species. Points in the panels at the right represent the beta diversity of species subsets that went locally extinct (orange) and colonized the sites (blue).

Regional-scale disturbance intensity can strongly determine the consequences of the local extinction and colonization processes for beta diversity (Caswell and Cohen 1991). For example, intense disturbance can make the regional environment more homogeneous, but moderate disturbance can create mosaics of different habitat types, some of which could act as refugial patches or lifeboats (Franklin et al. 1997, Gustafsson et al. 2012, Salo and Kouki 2018). These patches can prevent species from going locally extinct and can thereby increase beta diversity. Furthermore, such patches can serve as sources from which species recolonize the disturbed sites (Banks et al. 2011). However, such disturbance-intensity dependency of beta diversity, mediated by extinction and colonization processes, has received no investigation. With experiments manipulating disturbance intensity, one can test its effects on colonization–extinction dynamics and how this might translate into community homogenization or differentiation. Findings from such tests can help to better cope with future disturbance regimes and provide guidance for regional disturbance-based management.

In this study, we disentangled the roles of the extinction and colonization in shaping beta diversity along disturbance gradients. We did this by separately analyzing the post-disturbance beta diversity and the beta diversity of the subsets of species that went locally extinct and that newly colonized the given sites. As the study system, we selected a boreal forest in Finland, where we experimentally applied forest fire and tree harvesting with different intensities in a fully-crossed design. Forest fire is the dominant natural disturbance in many boreal and temperate regions. Harvesting was implemented as clearcutting and retention harvesting. Retention harvesting is a silvicultural system in which some portions of the original stand are left unlogged to maintain biodiversity and ecosystem functioning (Gustafsson et al., 2012; Lindenmayer et al., 2012). In our experiment, groups of trees were retained in patches, with the number of patches being manipulated. Fire and harvesting are known to drive disturbance interactions (Buma 2015); that is, the downed woody debris produced by harvesting can fuel fire and thereby cause synergetic impacts (Lindenmayer et al. 2009, Buma 2015). Our experiment thus provided an ideal system with which to unravel how the gains and losses of species generate spatiotemporal variation in species diversity across a wide range of disturbance intensities.

## METHODS

### Experimental design and data collection

We conducted a replicated stand-scale experiment in a boreal forest in eastern Finland (63°10’N, 30°40’E). Before the treatments, the experimental sites consisted of 150-year-old coniferous stands, which were dominated by Scots pine *(Pinus sylvestris* L.), with sporadic occurrences of Norway spruce *(Picea abies* [L.] Karst.) and two birch species *(Betula pendula* Roth and *Betula pubescens* Ehrh.). The average pre-treatment stand volume was 288 m^3^ ha^-1^ (SD = 71.1). The mean annual temperature is 2 °C, and the mean monthly temperature ranges from −12 °C in January to 16 °C in July. The mean annual precipitation is 500 to 800 mm, approximately half of which falls as snow.

Our experiment featured a 2 × 4 factorial design, with burnt and unburnt (control) for fire, and four harvesting levels: clear cut (0% retention), small-amount retention (10 m^3^ ha^-1^ or 3.5% of preharvest volume), large-amount retention (50 m^3^ ha^-1^ or 17.4% of preharvest volume), and no harvest (100% retention; control). In the two retention treatments, the retained trees were aggregated in either three (small-amount retention) or five (large-amount retention) groups of similar size. We selected 24 stands, each ca. 3 to 5 ha in size, located within a 20 km × 30 km area. Each stand was subjected to one of the eight treatment combinations, with three replicates for each combination. The stands were harvested in the winter of 2000–01 and burnt in the summer of 2001. See Hyvärinen et al. (2005) and Johnson et al. (2014) for further details of the experimental practices.

Field measurements followed the before-after control-impact (BACI) principle (Green 1979). In 2000, the summer before the treatments, we established fifteen 2 m × 2 m plots in each stand. The plots were placed systematically 20 m apart from each other in the center area of each stand. We recorded the presence and absence of all ground-living vascular plant, bryophyte, and lichen species in each plot. We considered these three groups to collectively represent the ground vegetation, as they often compete for similar resources and constitute a guild. Species were identified in the field or in the laboratory under a microscope. The nomenclature for vascular plants, bryophytes, and lichens followed Karlsson (1998), Hallingbäck et al. (2006), and Santesson et al. (2004), respectively. The same survey was repeated in 2003 (i.e., two years after treatment) and in 2011 (i.e., ten years after). The total sample size was *n* = 1080 (8 treatment combinations × 3 stands × 15 plots × 3 survey years).

### Statistical analyses

Beta diversity was defined as the extent of compositional dissimilarity within each stand. We used the Raup–Crick index (Raup and Crick 1979) to quantify the dissimilarity among local communities (= plots). There are a number of beta diversity metrics, and the choice of which to use depends on the research question (Anderson et al. 2011). We selected the Raup–Crick index because it controls for random sampling effects based on null models (Chase et al. 2011, Catano et al. 2017). Other beta diversity metrics such as the Whittaker, Jaccard, and Sørensen indices are known to be confounded by sampling effects that derive from variations in alpha and gamma diversity (Anderson et al. 2011). For example, if disturbance increases the mean alpha diversity and the gamma diversity remains unchanged, the observed beta diversity would necessarily decrease (e.g., β = γ/α) (Catano et al. 2017). This decrease in beta diversity would be indistinguishable from a pattern based on random sampling from the species pool and thus would not reflect selective homogenization (Chase et al. 2011, Catano et al. 2017). In this regard, the Raup–Crick index is capable of quantifying the compositional dissimilarity that is independent of the among-plot variation in alpha and gamma diversity (Vellend et al. 2007, Chase et al. 2011, Catano et al. 2017). We used the mean pairwise dissimilarity among the 15 plots to represent beta diversity in a given stand. The set of 15 plots was defined as the group within which species presence/absence was randomized.

Beta diversity was calculated for (i) species that were present in each survey year, (ii) species that went locally extinct within plots between two survey years (i.e., extinction beta diversity), and (iii) species that newly colonized each plot between two survey years (i.e., colonization beta diversity) (Fig. 1). We tested the effects of fire, harvesting, and their interaction on beta diversity using two-way ANOVA with beta error distributions and logit-link functions.

Alpha and gamma diversity was defined as the number of species in each plot and stand, respectively. We tested the effects of fire, harvesting, and their interaction on alpha diversity using two-way analysis of variance (ANOVA) with ‘stand’ as a random effect, and the effects on gamma diversity using two-way ANOVA. The tests were conducted for each survey year. We used Poisson error distributions and log-link functions for both diversity measures.

We defined the numbers of species that went extinct within plots and within stands between two survey years as the extinction alpha and gamma diversity, respectively. We defined the numbers of species that colonized each plot and each stand between two survey years as the colonization alpha and gamma diversity, respectively. The responses of these variables to fire, harvesting, and their interaction were tested using two-way ANOVA with ‘stand’ as the random effect for extinction and colonization alpha diversity, and two-way ANOVA for extinction and colonization gamma diversity. Poisson error distributions and log-link functions were used for both diversity measures.

We examined temporal species turnover as the compositional dissimilarity between the two survey years in each plot, which we quantified using the Jaccard and Sørensen indices. We did not use the Raup–Crick index here because we were focusing on the absolute extent of temporal dissimilarity, not the underlying assembly mechanisms. The Raup–Crick index also cannot quantify the dissimilarity when each community set, within which species co-occurrences are randomized, consists only of two samples (i.e., two surveys in different years). We used two indices (Jaccard and Sørensen) in order to confirm the robustness of our results. We tested the effects of fire, harvesting, and their interaction on temporal species turnover using two-way ANOVA with beta error distributions and logit-link functions. All analyses were performed using R 3.5.2 (R Core Team 2018).

## RESULTS

Alpha and gamma diversity did not differ significantly among the stands before the treatments (Fig. 2a, d), but changed dynamically after fire and harvesting. Burnt stands had significantly lower alpha and gamma diversity than the unburnt stands two years after the treatments (Fig. 2b, e). The fitted models showed that fire decreased the mean alpha diversity to 75% of the controls (i.e., 9.6 species in the burnt stands and 12.8 species in the unburnt stands, on average; Fig. 2b), and decreased the mean gamma diversity to 67% of the controls (i.e., 24.5 species in the burnt stands and 36.5 species in the unburnt stands; Fig. 2e). Ten years after the treatments, alpha and gamma diversity were no longer affected by fire, but were significantly higher in the harvested stands (clearcuts and retention stands) than in the unharvested stands (Fig. 2c, f). The mean alpha diversity in the harvested stands was 1.37 to 1.46 times that in the unharvested stands (Fig. 2c), and mean gamma diversity was 1.38 to 1.47 times that in the unharvested stands (Fig. 2f).

**Fig. 2.**
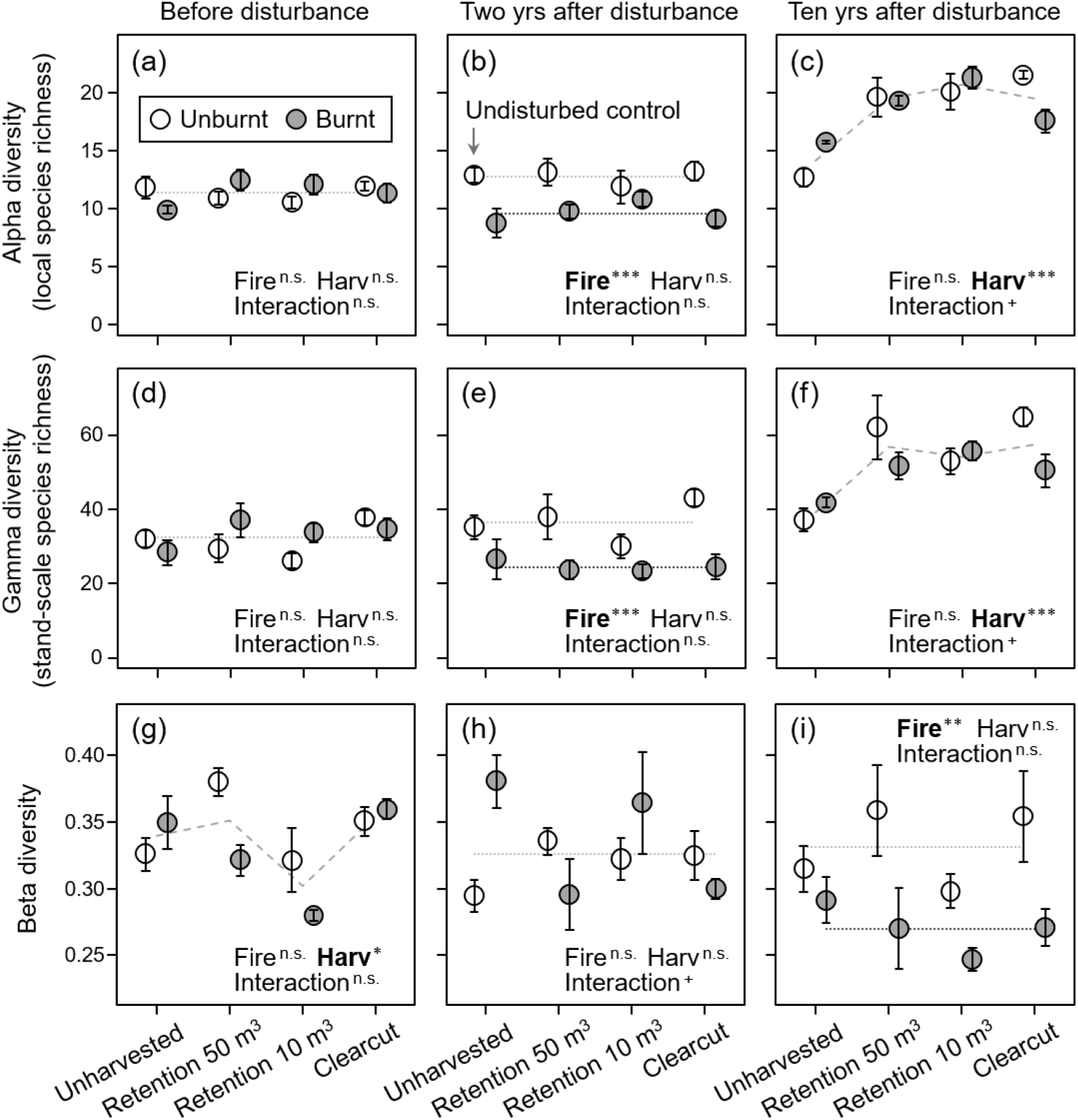
Effects of fire and harvesting disturbance on (a–c) alpha, (d–f) gamma, and (g–i) beta diversity. “Retention 50 m^3^” and “10 m^3^” indicate harvesting treatments in which those volumes of trees per hectare were left unlogged. The results of two-way ANOVA are shown in each panel: Harv = harvesting; Interaction = fire × harvesting. Significance: *** *P* < 0.001; ** *P* < 0.01; * *P* < 0.05; + *P* < 0.10; n.s., *P* ≥ 0.10. Variables are shown in boldface when their effects were significant (*P* < 0.05). Lines represent the fitted models for significant variables. Black lines and dashed lines indicate significant effects of fire and harvesting, respectively. Values are means ± SE.

Beta diversity showed different patterns depending on the survey years (Fig. 2g, h, i). Before the treatments, beta diversity differed among the stands subjected to different harvesting intensities (Fig. 2g). Two years after the treatments, this difference became undetectable, and fire also showed no effect on beta diversity (Fig. 2h). Ten years after the treatments, beta diversity showed no significant difference among the harvesting levels, but became significantly lower in the burnt stands than in the unburnt stands (Fig. 2i).

Extinction beta diversity at the first survey interval (i.e., beta diversity of the sets of species that went locally extinct between pre-treatment and two years after the treatments) was lower in the disturbed stands than in the undisturbed stands (Fig. 3a). This pattern was particularly pronounced when fire and harvesting were combined (although the interaction effect [fire × harvesting] was not significant); mean β ranged from 0.40 to 0.46 in the burnt and harvested stands, versus 0.93 in the undisturbed stands (Fig. 3a). Between two and ten years after the treatments, only harvesting had a significant effect on extinction beta diversity (Fig. 3b). The decreased extinction beta diversity at both survey intervals indicates that disturbance excluded similar suites of species across the plots within each stand (i.e., case 2 in Fig. 1).

**Fig. 3.**
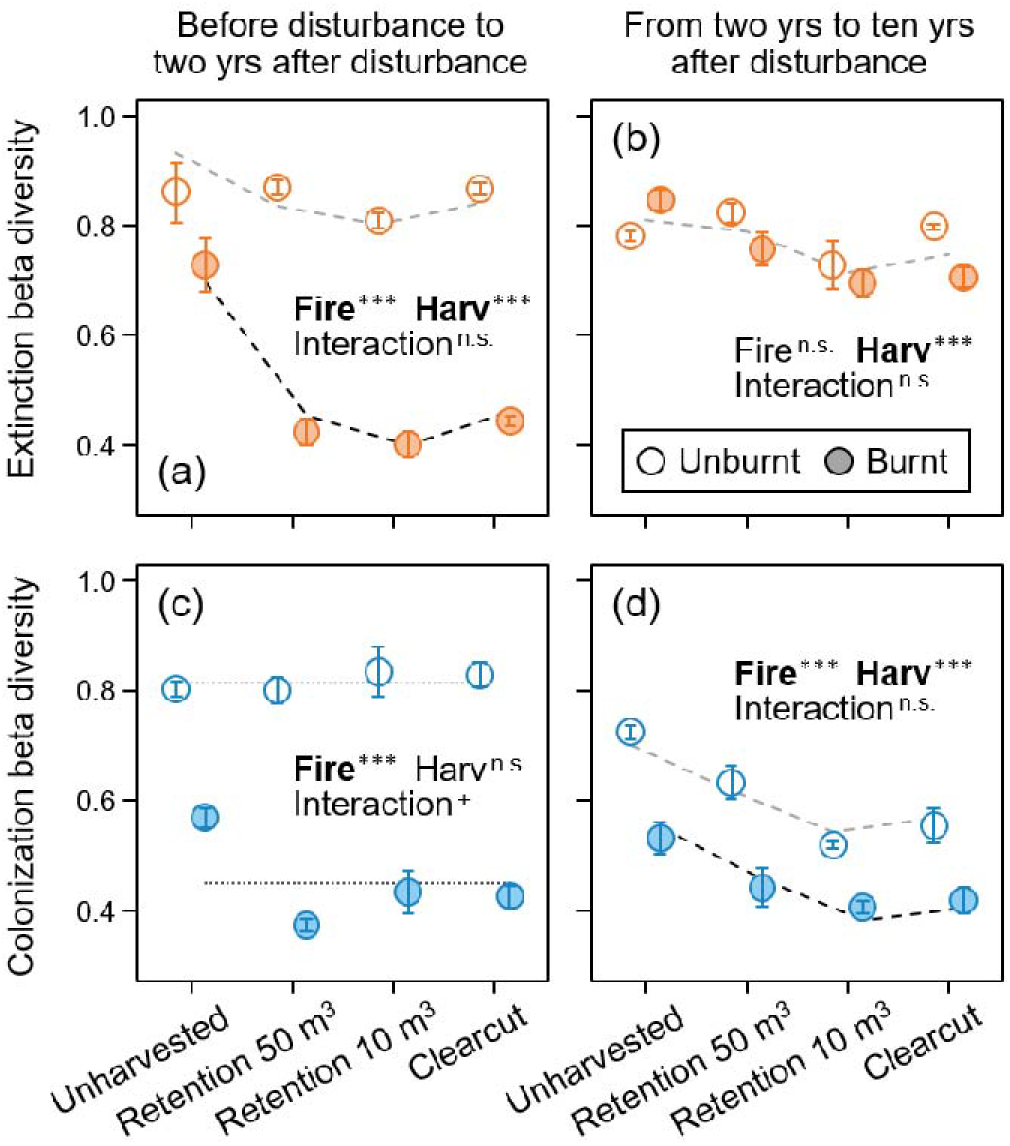
Effects of fire and harvesting disturbance on beta diversity of (a, b) species that went locally extinct within plots and (c, d) species that newly colonized the plots. “Retention 50 m^3^” and “10 m^3^” indicate harvesting treatments in which those volumes of trees per hectare were left unlogged. The results of two-way ANOVA are shown in each panel: Harv = harvesting; Interaction = fire × harvesting. Significance: \*\*\**P* < 0.001; \**P* < 0.05; +*P* < 0.10; n.s., *P* ≥ 0.10. Variables are shown in boldface when their effects were significant (*P* < 0.05). Lines represent the fitted models for significant variables. Black lines and dashed lines indicate significant effects of fire and harvesting, respectively. Values are means ± SE.

Colonization beta diversity at the first survey interval (i.e., beta diversity of the set of species that newly colonized the plots between pre-treatment and two years after the treatments) was lower in the burnt stands than in the unburnt stands (Fig. 3c). The interaction effect (fire × harvesting) had a marginally significant effect (*P* < 0.10), indicating the potential for synergetic negative impacts of fire and harvesting on colonization beta diversity (Fig. 3c). Fire and harvesting reduced the colonization beta diversity from two to ten years after the treatments (Fig. 3d). The decreased colonization beta diversity at both survey intervals indicates that similar sets of species have colonized different plots within each stand after disturbance (i.e., case 4 in Fig. 1). Extinction and colonization alpha and gamma diversity were significantly influenced by fire and harvesting (Fig. S1).

The rates of temporal species turnover were increased significantly by both fire and harvesting (Fig. S2). The Jaccard (Fig. S2a, b) and Sørensen indices (Fig. S2c, d) showed the same qualitative pattern. Between pre-treatment and two years after the treatments, the temporal species turnovers were large in the burnt stands (Fig. S2a, c). From two to ten years after the treatments, fire continued to be the dominant driver of species turnover, yet the positive effects of harvesting increased with increasing intensity (Figs. S2b, d).

## DISCUSSION

As is common in community assembly research, studies on beta diversity have typically overlooked the dynamic nature it inherently bears (Fukami and Nakajima 2011, Li et al. 2016, Tatsumi et al. 2019). In fact, though ample evidence exists for both cases in which disturbance increased or decreased beta diversity, the dynamic processes that underlie those patterns had not necessarily been articulated. Here, we proposed a conceptual foundation to disentangle the extinction and colonization components of beta diversity. Specifically, we defined six processes by which colonization–extinction dynamics can drive community homogenization or differentiation (Fig. 1). Analyses based on this concept allowed us to detect nonrandom dis- and re-assembly dynamics in plant communities.

The difference in beta diversity was undetectable across an experimental disturbance-intensity gradient two years after the treatments (Fig. 2h). From this result, we might conclude that disturbance caused no impact on how species assemble across space. Nevertheless, when we analyzed the extinction and colonization beta diversity (i.e., beta diversity of the sets of species that went locally extinct and that newly colonized the given sites, respectively), we found that both were significantly lower in the disturbed stands than in the undisturbed controls (Fig. 3a, c). These patterns indicate that the disturbance excluded similar species subsets across plots, making communities differentiate (case 2 in Fig. 1), but at the same time induced spatially uniform colonization of new species, causing communities to become more homogeneous (case 4 in Fig. 1). Consequently, the effects of these two processes canceled each other out (Fig. 2h). We further found that beta diversity decreased in the burnt stands relative to the unburnt stands after ten years (Fig. 2i). This pattern was explained by the fact that the spatially uniform colonization continued to make the communities become more homogeneous over time (case 4 in Fig. 1; Fig. 3d). Overall, our results provide evidence that the beta diversity at our experimental site was determined by nonrandom extinction–colonization dynamics and, moreover, that the relative importance of extinction and colonization components of beta diversity changed with time after disturbance.

### Extinction and colonization beta diversity driven by disturbance

Extinction beta diversity was lower in the burnt stands than in the unburnt stands two years after the treatments (Fig. 3a). This result indicates that the fire disproportionately excluded habitat generalist species that commonly occurred across the given stand (case 2 in Fig. 1). It is important to note here that we quantified beta diversity using the Raup–Crick index, which corrects for random sampling effects (Vellend et al. 2007, Chase et al. 2011). That is, our finding that the fire reduced extinction beta diversity (Fig. 3a) does not result simply from the fact that generalists had high local-extinction frequencies due to their widespread occurrence, but rather indicates that the fire selectively excluded generalists over specialists. Tolerance to fire is a cost-intensive plant trait that requires considerable structural and energy investments (Wahid et al. 2007). Having this trait might thus come at the expense of species to have narrow niche widths, resulting in fire to cause more severe damage on habitat generalists than on specialists. Furthermore, we found that the effects of fire on extinction beta diversity became undetectable between two and ten years after the treatments (Fig. 3b). This result further suggests that the direct effects of burning influenced extinction beta diversity more strongly than subsequent environmental alterations, such as a post-fire increase in soil pH (Čugunovs et al. 2017).

Harvesting, on the other hand, continued to reduce extinction beta diversity until ten years after the treatments (Fig. 3 a, b). This result suggests prolonged extinction of generalists that were common before harvesting. We can see such case, in which beta diversity continues to change owing to time-delayed but deterministic extinctions, as an example of an ecosystem incurring extinction debts of beta diversity. Extinction debt, which was originally coined to refer to the number of species that are expected to eventually go extinct, occurs because of delays in population responses to environmental changes (Tilman et al. 1994). In our harvested stands, the extinction debt of beta diversity occurred likely because some populations of shade-adapted species were able to persist after the disturbance (≤2 years), despite the stresses caused by increased light and competition with open-habitat species. In human-altered ecosystems, it can take more than a century to pay off an extinction debt (Vellend et al. 2006). Our results indicate that spatially replicated, long-term monitoring is necessary to understand the potential long-lasting effects of disturbance on the future spatial structure of ecological communities.

Colonization beta diversity decreased in stands disturbed by fire and/or harvesting (Fig. 3c, d). This suggests that similar suites of species, likely represented by long-distance dispersers and post-disturbance germinators/resprouters (Hollingsworth et al. 2013, Johnson et al. 2014), uniformly covered the stands after disturbance (case 4 in Fig. 1). Until two years after the treatments, colonization beta diversity was reduced especially in stands where fire and harvesting were both applied (*P* < 0.10; Fig. 3c), indicating the signature of disturbance interaction. In fact, at our experimental site, fire intensities (measured as the post-fire humus depth and height of charred bark) were significantly higher in the harvested stands than in the unharvested stands (Hyvärinen et al. 2005). Thus, the low colonization beta diversity in the burnt and harvested stands (Fig. 3c) suggests that large fires, fueled by the downed woody debris, have homogenized the environment and made spaces available for new species to colonize across the stands.

Furthermore, we found that the large-amount retention stands had higher colonization beta diversity ten years after the treatment than the small-amount retention stands and clearcuts (Fig. 3d). This pattern suggests that the retention patches served as refugia (Franklin et al. 1997; Gustafsson et al. 2012), which allowed species with poor dispersal abilities, that otherwise could have hardly reached the post-disturbance areas, to recolonize them sporadically. In future studies, it would be worthwhile exploring whether such “chance colonization” will amplify community differentiation over time (Chase 2007; Fukami & Nakajima 2011).

### General and future applications

The concept of extinction and colonization beta diversity (Fig. 1) can be applied to any taxonomic group and only requires a minimum of two community data taken at different points in time. It is applicable not only in the context of ecosystem disturbance, but also of any environmental changes that can drive spatiotemporal species turnover (e.g., climate change, biological invasions, habitat fragmentation, or ecosystem restoration). For example, extinction and colonization beta diversity can be utilized in a restoration project that aims to reverse biotic homogenization — it can be used to identify whether a given recovery of beta diversity is due to local species extinctions (case 2 or 3 in Fig. 1) or (re)colonization of new species (Case 4), the latter of which may often be the ultimate goal in restoration.

Extinction and colonization beta diversity can also be highly relevant for understanding ecosystem functionality. Recent studies using snapshot data have shown that spatial dissimilarity in species composition can affect regional-scale ecosystem functioning as strongly as, if not more strongly than, local species richness (van der Plas et al. 2016, Hautier et al. 2018, Mori et al. 2018). By linking extinction and colonization beta diversity with temporal changes in ecosystem functionality, one could directly discern what species subsets should be removed or added to enhance ecosystem functioning. Moreover, it can be possible to quantify the spatial variation in the capability of community members to withstand extinction and to recolonize the given sites during and after environmental perturbations (i.e., resistance and resilience of species composition; *sensu* Allison and Martiny 2008), and test how this might affect the stability of regional-scale ecosystem functioning (Loreau et al. 2003). This can potentially be done using our framework and expanding it to account for species’ abundances and functional traits, which are often associated with ecosystem functioning (Hooper et al. 2005, Cadotte 2017).

## Conclusions

In this study, we defined six processes through which species extinction and colonization shape beta diversity. Analyses based on repeated community measurements revealed that relying only on snapshot data can sometimes mislead us to superficial perceptions that disturbance has not caused any detectable variation. Our results indicated that accounting for colonization–extinction dynamics could help us test and expand some current ideas in community ecology. Specifically, extinction beta diversity allowed us to detect nonrandom extinctions, where a higher proportion of habitat generalists compared to specialists was removed. We also found time-delayed changes in spatial community variation, which can be seen as a novel type of extinction debt. Colonization beta diversity indicated that retained habitats could indeed serve as refugia from which species recolonize a disturbed site, providing one of the first evidence for the conservation benefits of retention patches at the community level. Overall, our results show that the concepts of extinction and colonization beta diversity form a useful link between community assembly studies, which have often been spatially framed, and ecological disturbance and dynamics studies, which have focused primarily on temporal changes. Dynamic appraisals of beta diversity will help us to better understand the spatiotemporal organization of biodiversity and its consequences for ecosystem functioning.

## ACKNOWLEDGEMENTS

We thank Mervi Hakulinen, Hanna Keski-Karhu, Anneli Uotila, Samuel Johnson, Osmo Heikkala, and Mikko Heikura for their assistance during the fieldwork. We are also grateful to Akira S. Mori, Ryo Kitagawa, Åsa Ranlund, and Lena Gustafsson for helpful discussions about the manuscript. We thank Bill Liu for English language editing. ST was supported by a JSPS Overseas Research Fellowship (No. 201860500) and Grants-in-Aid for JSPS Research Fellows PD (No. 15J10614) and for Young Scientists B (No. 16K18715) from the Japan Society for the Promotion of Science. JK received funding for a sabbatical from the Maj and Tor Nessling Foundation during the preparation of this manuscript.

