## SupplementaryMaterials for "Partitioning the colonization and extinction components of beta diversity: Spatiotemporal species turnover across disturbance gradients"

### Supporting Information

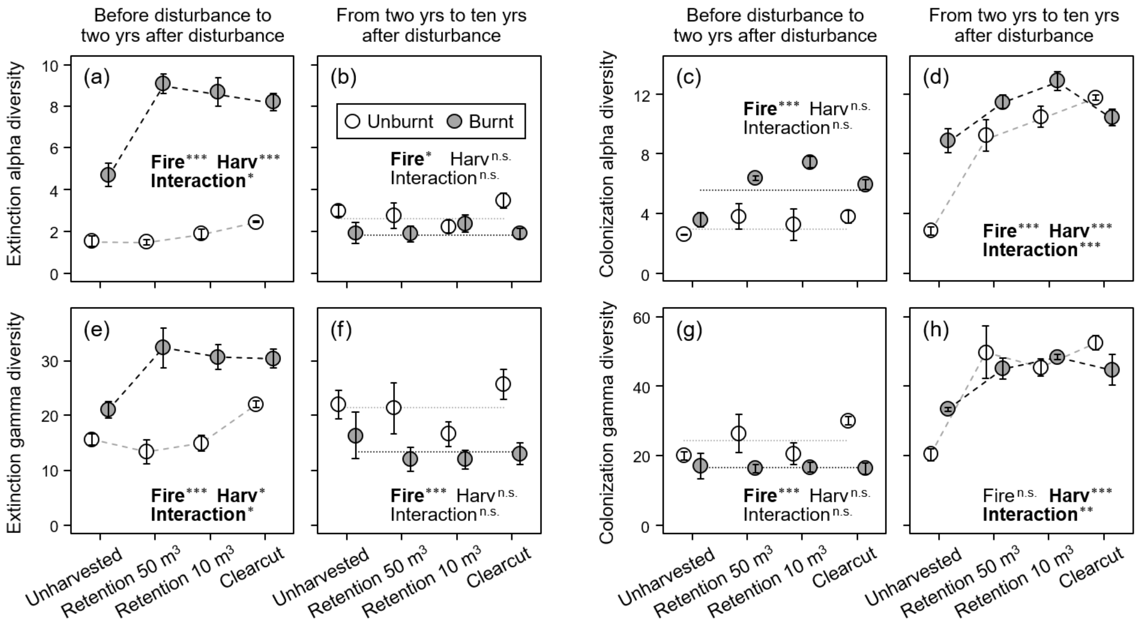


**Fig. S1** Effects of fire and harvesting disturbance on the numbers of (a, b) species that went locally extinct within plots, (c, d) species that newly colonized each plot, (e, f) species that went extinct within stands, and (g, h) species that newly colonized each stand. "Retention 50 $m^3$" and "10 $m^3$" indicate harvesting treatments in which those volumes of trees per hectare were left unlogged. The results of two-way ANOVA are shown in each panel: Harv = harvesting; Interaction = fire × harvesting. Significance: *** $P < 0.001$; ** $P < 0.01$; * $P < 0.05$; n.s., $P \geq 0.05$. Variables are shown in boldface when their effects were significant ($P < 0.05$). Lines represent the fitted models for significant variables. Values are means ± SE.

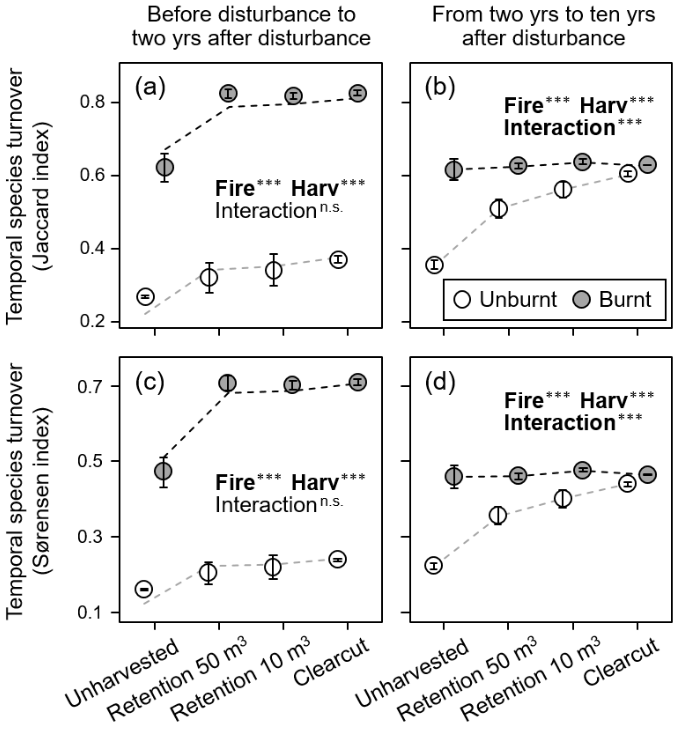


**Fig. S2** Effects of fire and harvesting disturbance on temporal species turnover as measured by (a, b) the Jaccard index and (c, d) the Sørensen index. "Retention 50 m$^3$" and "10 m$^3$" indicate harvesting treatments in which those volumes of trees per hectare were left unlogged. The results of two-way ANOVA are shown in each panel: Harv = harvesting; Interaction = fire × harvesting. Significance: *** $P < 0.001$; n.s., $P \geq 0.05$. Variables are shown in boldface when their effects were significant ($P < 0.05$). Lines represent the fitted models for significant variables. Values are means ± SE.
